# Within-host succession from an OXA-48-like to an NDM-type carbapenemase in clonal ST361 *Escherichia coli* recovered from sequential urinary and bloodstream infection

**DOI:** 10.64898/2026.07.27.741121

**Authors:** Leon Kassabian, Charbel Al Khoury, George F. Araj, Sima Tokajian

## Abstract

Carbapenem-resistant *Escherichia coli* (CREc) recovered sequentially from one patient typically retain the same carbapenemase, with escalating resistance usually attributed to porin loss combined with pre-existing β-lactamase expression. We used whole-genome sequencing to characterize a clonal pair of CREc isolates, CAEC145 and CAEC155, recovered 25 days apart from a hospitalized patient with sequential urinary and bloodstream infection. Both belonged to sequence type 361 (ST361), phylogroup A, serotype O-nontypeable:H30, and were separated by only 28 core-genome SNPs, confirming clonal relatedness. Despite this, the isolates differed sharply in carbapenemase content. CAEC145 carried *bla*_OXA-1207_, a recently described OXA-48-family variant, on a conjugative IncFII(pCoo)/ColKP3 plasmid, whereas CAEC155 lacked this determinant and instead harbored *bla*_NDM-4_ on a conserved IncX3 plasmid nearly identical to pJEG027, a member of a globally disseminated IncX3 lineage. This genotypic shift tracked a clear phenotypic transition. CAEC145 remained susceptible to imipenem and meropenem while resistant to ertapenem, whereas CAEC155 showed uniform high-level resistance to all three carbapenems and to ceftazidime-avibactam. Both isolates, however, remained susceptible to imipenem-relebactam, meropenem-vaborbactam, and cefiderocol. Comparative genomics linked the *bla*_OXA-1207_ element to a ΔTn*6361* transposon structure also found in the original German isolates where *bla*_OXA-1207_ was first described, and a 137-genome core-genome phylogeny placed both isolates within a globally disseminated ST361 lineage carrying multiple carbapenemase classes. These findings document, to our knowledge, the first within-host succession from an OXA-48-like to an NDM-type carbapenemase during a single sequential *E. coli* infection, driven by plasmid-level displacement rather than in-place gene evolution, with implications for genomic surveillance and antibiotic selection.

## Introduction

Carbapenem-resistant Enterobacterales (CRE) remain a major source of difficult-to-treat infection worldwide. Limited therapeutic options and their capacity for rapid dissemination led the World Health Organization to rank them as critical in its most recent Bacterial Priority Pathogens List, with comparable trends reported in Lebanon (1–3). *Escherichia coli* is among the most clinically significant species in this group. It is a near-universal member of the gut microbiota, yet given appropriate virulence determinants and host conditions it is also a leading cause of invasive and extraintestinal disease. This dual role allows resistant lineages to persist as a colonizing reservoir between episodes of overt infection (4). Certain *E. coli* lineages, including pandemic high-risk clones that have driven the global spread of carbapenem resistance, combine this reservoir capacity with a tendency toward recurrent or relapsing infection in the same host (5). This creates a prolonged within-host setting in which the infecting lineage faces repeated antibiotic selection, favoring ongoing genomic and plasmid-level evolution (5).

The clinical stakes are magnified by the diversity of carbapenemases circulating in Enterobacterales. OXA-48-like class D serine β-lactamases are now endemic in many regions, yet their impact is readily underestimated. They hydrolyze carbapenems weakly and inconsistently, show little activity against extended-spectrum cephalosporins, and can escape routine phenotypic screening despite conferring clinically relevant carbapenem resistance (6). In Lebanon, OXA-48-family carbapenemases were first documented in a clinical *E. coli* isolate in 2018, and have since become part of the local resistance landscape (7).

NDM-family metallo-β-lactamases represent the opposite extreme. As zinc-dependent class B enzymes, they hydrolyze nearly the entire β-lactam class with high efficiency, sparing only aztreonam (8). They are also refractory to the diazabicyclooctane and boronate inhibitors (avibactam, relebactam, and vaborbactam) that have restored the activity of several carbapenems and cephalosporins against serine-carbapenemase producers (9). Aztreonam-avibactam remains one of the few reliable options against NDM-producing Enterobacterales, yet resistance has already been reported in both food-chain and clinical settings (10, 11). A within-host transition from an OXA-48-like determinant to an NDM variant therefore represents a change in carbapenemase class, not an incremental worsening of resistance, with direct consequences for empirical and definitive antibiotic selection.

*bla*_OXA-1207_, a recently characterized OXA-48 family-variant was first described by Sommer et al. (2025) in two epidemiologically unrelated *E. coli* isolates (ST405, ST4405) from a German university hospital (12). OXA-1207 differs from OXA-181 by two substitutions, 214G and 244W. It was found on a self-transmissible plasmid bearing a ΔTn*6361* transposon structure derived from IncX3-associated OXA-181 lineages, with a conjugation frequency (mean 1.2 × 10⁻¹) substantially exceeding rates typical of OXA-48-family plasmids (12). Suleiman et al. (2025) subsequently reported OXA-1207 in an ST617 *E. coli* isolate from a pediatric patient in Qatar, in a genetic context containing IS*Kpn19* transposase and Tn*3* as previously reported by Sommer et al. (2025) (12, 13). Whether *bla*_OXA-1207_ persists, is lost, or is succeeded by a carbapenemase of a different class over a single relapsing infection has not previously been documented.

The molecular basis for such a within-host succession lies in plasmid population dynamics under sustained antibiotic selection. Plasmids are classified into incompatibility (Inc) groups by replicon typing (14). Plasmids sharing replication or partitioning machinery generally cannot stably co-reside in the same cell, whereas plasmids of different Inc groups can be maintained together (15). Independently of compatibility, plasmid carriage imposes a metabolic burden that reduces host fitness relative to plasmid-free or differently plasmid-bearing competitors (16). These dynamics are consistent with the acquisition of *bla*_NDM-4_, a single-substitution (M154L) variant of *bla*_NDM-1_ with increased hydrolytic activity, first reported in *E. coli* from India (17). NDM determinants are frequently located adjacent to a bleomycin-resistance cassette on the highly stable, conjugation-proficient IncX3 replicon, which has driven much of the global spread of NDM (18, 19).

Here, we report the whole-genome characterization of paired *E. coli* isolates, CAEC145 and CAEC155, recovered from the same patient during a sequential infection. Core-genome MLST (cgMLST) and pairwise SNP analysis confirmed that the isolates were clonally related, yet they differ sharply in carbapenemase content: CAEC145 carries *bla*_OXA-1207_ on a conjugative plasmid, whereas CAEC155 lacks this determinant entirely and instead carries *bla*_NDM-4_ co-located with *ble* on an IncX3 plasmid. We used plasmid replicon typing and mobile-element reconstruction to map the genetic context underlying this apparent in vivo carbapenemase class switch, place the isolate pair within the global ST361 lineage using a 137-genome core-genome phylogeny, and describe what is, to our knowledge, the first documented within-host succession from an OXA-48-like to an NDM-type carbapenemase during a sequential *E. coli* infection.

## Methods

### Bacterial isolates and identification

The two isolates were recovered from urine and blood specimens submitted sequentially by the same hospitalized patient for culture at the Clinical Microbiology Laboratory, Department of Pathology and Laboratory Medicine, American University of Beirut Medical Center, in April and May 2025, respectively. Species identification was performed by matrix-assisted laser desorption ionization–time of flight mass spectrometry (MALDI-TOF MS) using the MALDI Biotyper system with accompanying software (Bruker Daltonics, Bremen, Germany).

### Antimicrobial susceptibility testing and carbapenemase detection

Screening for carbapenem resistance was performed by disk diffusion (DD) using an ertapenem disk (10 µg) on Mueller-Hinton agar (MHA) per CLSI M100 (35th ed.) (20), incubated at 37 °C for 18 h. Confirmatory MICs for ertapenem, imipenem, meropenem, and ceftazidime-avibactam were determined by the E-test method (AB BIODISK, Solna, Sweden) as previously described by Araj et al. (2). Routine susceptibility testing by Disk Diffusion was performed on MHA using commercial disks (BBL, Becton Dickinson, USA) for the following agents: amikacin (30 µg), gentamicin (10 µg), trimethoprim-sulfamethoxazole (1.25/23.75 µg), tigecycline (15 µg), fosfomycin/trometamol (200 µg), aztreonam (30 µg), ceftazidime (30 µg), cefepime (30 µg), and levofloxacin (5 µg). Susceptibility to the novel β-lactam/β-lactamase inhibitor combinations imipenem-relebactam and meropenem-vaborbactam, and to cefiderocol, was also assessed by DD. All zone diameters were interpreted per CLSI M100 (35th ed.) (20), with the following exceptions: tigecycline lacks a CLSI interpretive breakpoint for Enterobacterales and zone diameters are reported without categorical interpretation; for cefepime, CLSI M100 suppression rules were applied, whereby susceptible and susceptible-dose-dependent results are reported as Resistant for confirmed carbapenemase-producing isolates. Quality control was performed in parallel using *E. coli* ATCC 25922 and *Pseudomonas aeruginosa* ATCC 27853.

Phenotypic carbapenemase detection was performed using the NG-Test CARBA-5 multiplex immunochromatographic assay (NG Biotech, Guipry, France), which enables simultaneous detection and differentiation of the five most prevalent carbapenemase families: KPC, OXA-48-like, VIM, IMP, and NDM. The assay was performed and interpreted according to the manufacturer’s instructions.

### DNA extraction and whole-genome sequencing

Genomic DNA was extracted from overnight cultures using the Sigma-Aldrich DNA extraction kit (Sigma-Aldrich; St. Louis, USA) per the manufacturer’s instructions. Short-read libraries were constructed using the Nextera XT DNA Library Preparation Kit (Illumina) and sequenced on an Illumina MiSeq with a paired-end 500-cycle protocol (2 × 250 bp). Long-read libraries were prepared using the ONT Rapid Barcoding Kit V14 (SQK-RBK114.24/96); end-prep and barcode ligation were performed on 100–200 fmol of sample in 65 µL nuclease-free water, followed by adapter ligation and cleanup with NEB ligase and Agencourt AMPure XP beads (Beckman Coulter, USA), yielding a final adapter-ligated library of 50–100 fmol. The library was loaded onto an R10.4.1 flow cell and sequenced on a MinION device (Oxford Nanopore Technologies).

### Genome assembly, annotation and analysis

Illumina raw reads were quality-trimmed and adapter-stripped with fastp (v1.3.1) (21). ONT raw data underwent basecalling with Guppy (v4.5.2) at a minimum Q-score of 9 using the high-accuracy model (dna_r9.4.1_450bps_hac.cfg; --num_callers 4, --cpu_threads_per_caller 4), with demultiplexing and barcode trimming performed using EXP-NBD104 and EXP-NBD114 kit configurations. Hybrid assemblies were generated with Unicycler (v0.5.1) (22) and assessed for quality and completeness with QUAST (v5.3.0) (23). Genome annotation was performed with the Bakta pipeline (v1.11.4) against the v6.0 light database (24).

Assembled genomes were profiled for virulence factors using ABRicate (v1.2.0) against the VFDB database (25), and for antimicrobial resistance (AMR) genes using AMRFinderPlus (v4.2.5, database v2025-12-03.1) (26). Plasmid replicons were identified with PlasmidFinder (v2.1.6), phylogroups assigned with ClermonTyping (27), and sequence types determined with mlst (v2.35.0; Seemann) using the Achtman 7-gene scheme (EnteroBase/PubMLST). Serotyping was performed with ECtyper (v1.0) (28).

### Genetic environment of resistance genes

The genetic environments of *bla*_OXA-1207_ (CAEC145) and *bla*_NDM-4_ (CAEC155) were compared against the German clinical isolates EC8729 (GCA_965650475) and EC8533 (GCA_965650485) and reference plasmids pKP3-A (JN205800), pIMP-Z1058 (KU051709), pMR3-OXA-181 (KM660724), pNDM-HN380 (JX104760), and pJEG027 (KM400601). Pairwise alignments were generated with minimap2 (v2.31, -x asm5) (29), and synteny maps were visualized using gggenomes (v1.1.3) in R. Alignment identity was calculated as matching bases divided by alignment block length, multiplied by 100.

### Core-genome phylogenetic analysis

Public ST361 *E. coli* genome assemblies were retrieved from NCBI. Lebanese isolates lacking deposited assemblies were assembled from raw SRA reads with Shovill v1.4.1 (https://github.com/tseemann/shovill). A stratified subset of 137 isolates representing global geographic diversity, with all Lebanese isolates retained, was selected for phylogenetic analysis (Material S1). Core-genome SNP calling was performed with Snippy v4.6.0 in contig mode (--ctgs) against a high-quality ST361 reference genome (SAMN53177129; GCA_057961095.1), and a core alignment was generated with snippy-core (https://github.com/tseemann/snippy). Recombinant regions were identified and masked with Gubbins (30), and a maximum-likelihood phylogeny was inferred from the recombination-filtered alignment with IQ-TREE2 using ModelFinder Plus (-m MFP) and 1,000 ultrafast bootstrap replicates. Pairwise SNP distances were calculated from the recombination-filtered alignment with snp-dists v1.2.0 (https://github.com/tseemann/snp-dists). The phylogeny was visualized as a circular cladogram with ggtree in R.

## Results

### Strain typing and clonal relatedness

CAEC145 and CAEC155 were both assigned to Clermont phylogroup A, sequence type 361 in the Achtman scheme and 650 in the Pasteur scheme, and serotype O-nontypeable:H30, with no O antigen assignable in silico for either isolate. The two isolates differed by 28 core-genome SNPs.

### Genome features, resistome, and virulome

The CAEC145 and CAEC155 assemblies totaled 5,168,932 bp and 5,261,684 bp, respectively, with GC contents of 50.67% and 50.72%. A total of 17 distinct AMR genes were detected in CAEC145 and 20 in CAEC155. A total of 111 and 115 distinct virulence genes were identified in CAEC145 and CAEC155, respectively (Table 1). Both isolates shared *bla*_EC_, *bla*_TEM-1_, *bla*_CMY-145_, and a conserved class 1 integron cassette array (*sul1*-*aadA5*-*dfrA17*-*qacEΔ1*). CAEC145 carried *bla*_OXA-1207_ as its sole carbapenemase; this gene was absent from CAEC155, which instead carried *bla*_NDM-4_, co-located with the bleomycin-resistance gene *ble*. CAEC155 additionally carried a second *bla*_TEM-1_ copy, *fosA4*, *tet(*M), second copies of *tet(*A), *aph(3’’)-Ib, aph*(*6*)*-Id*, and *floR* (Supplementary Material S2), and the complete salmochelin locus *iroBCDEN*, none of which were present in CAEC145 (Supplementary Material S3).

**Table 1.**
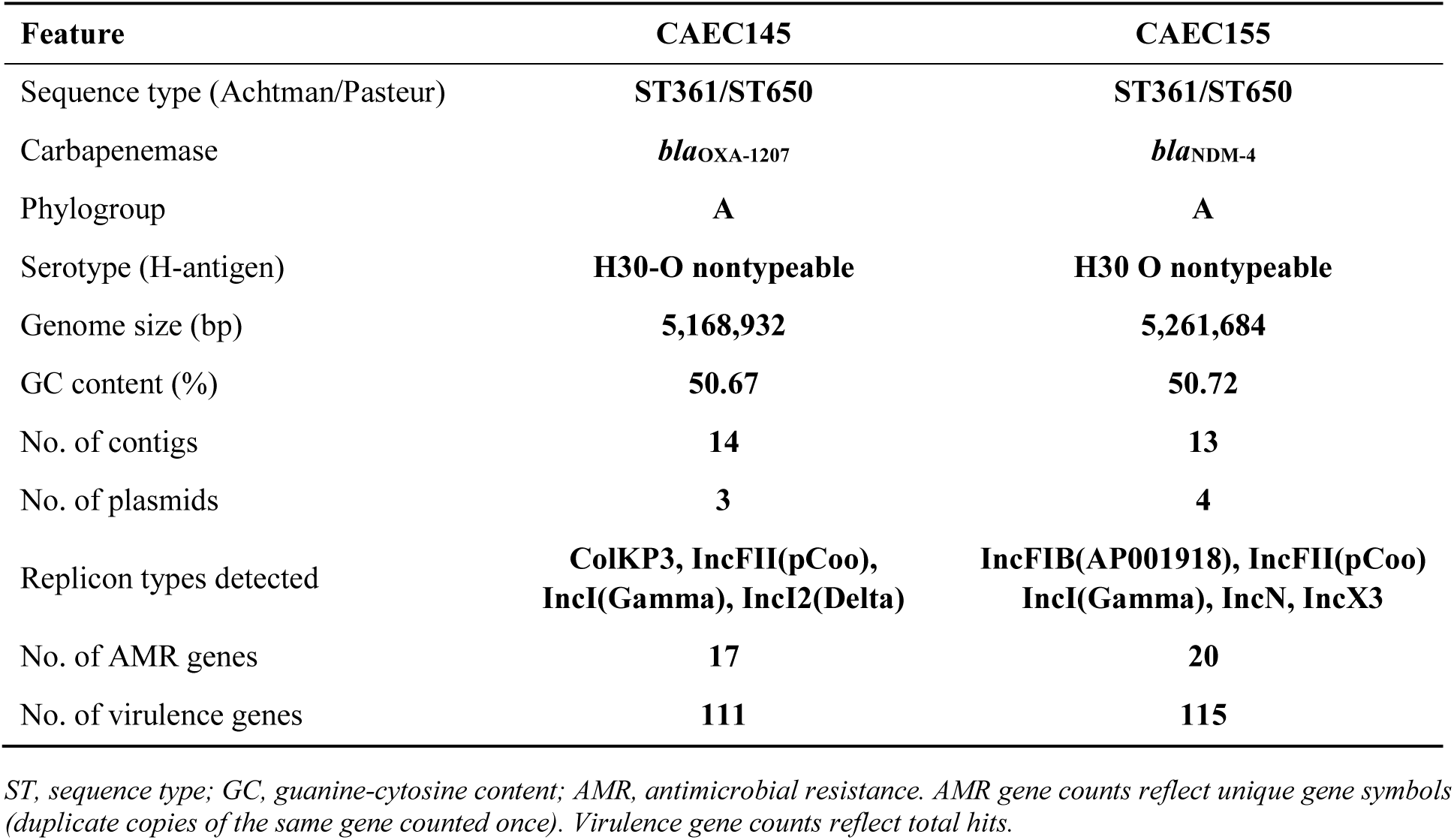
Genomic characteristics of the sequential urinary (CAEC145) and bloodstream (CAEC155) ST361 *Escherichia coli* isolates.

### Plasmid content and replicon typing

PlasmidFinder detected four replicons in CAEC145 (ColKP3, IncFII(pCoo), IncI(Gamma), IncI2(Delta)). The 138,706 bp IncFII(pCoo)/ColKP3 plasmid carried *bla*_OXA-1207_, *bla*_TEM-1_, *bla*_CMY-145_, *mph(A)*, and *qnrS1*. PlasmidFinder detected five replicons in CAEC155 (IncFIB(AP001918), IncFII(pCoo), IncI(Gamma), IncN, IncX3). These included a predicted IncFIA/IncFIB plasmid carrying the *iro* locus and a 57,003 bp IncI(Gamma)/K1 plasmid identical by MASH distance to that of CAEC145. The *bla*_NDM-4_-bearing contig was typed as IncX3 (100% identity, 374/374 bp), and an IS*Kox3* insertion element was detected in the genetic environment of *bla*_NDM-4_.

#### NDM genetic environment

The *bla*_NDM-4_ locus of CAEC155 shared 99.78% nucleotide identity with pJEG027 (IncX3-*bla*_NDM-4_) across the IS*5*-*bla*_NDM_-*ble*-*trpF*-*dsbD* cassette, and >99% identity with the prototype pNDM-HN380 (IncX3-*bla*_NDM-1_) (Figure 1). CAEC155 shared 99.87% nucleotide identity with pJEG027 (IncX3-blaNDM-4) across the complete plasmid backbone (45,280/45,337 bp; 99.97% query coverage)

**Figure 1.**
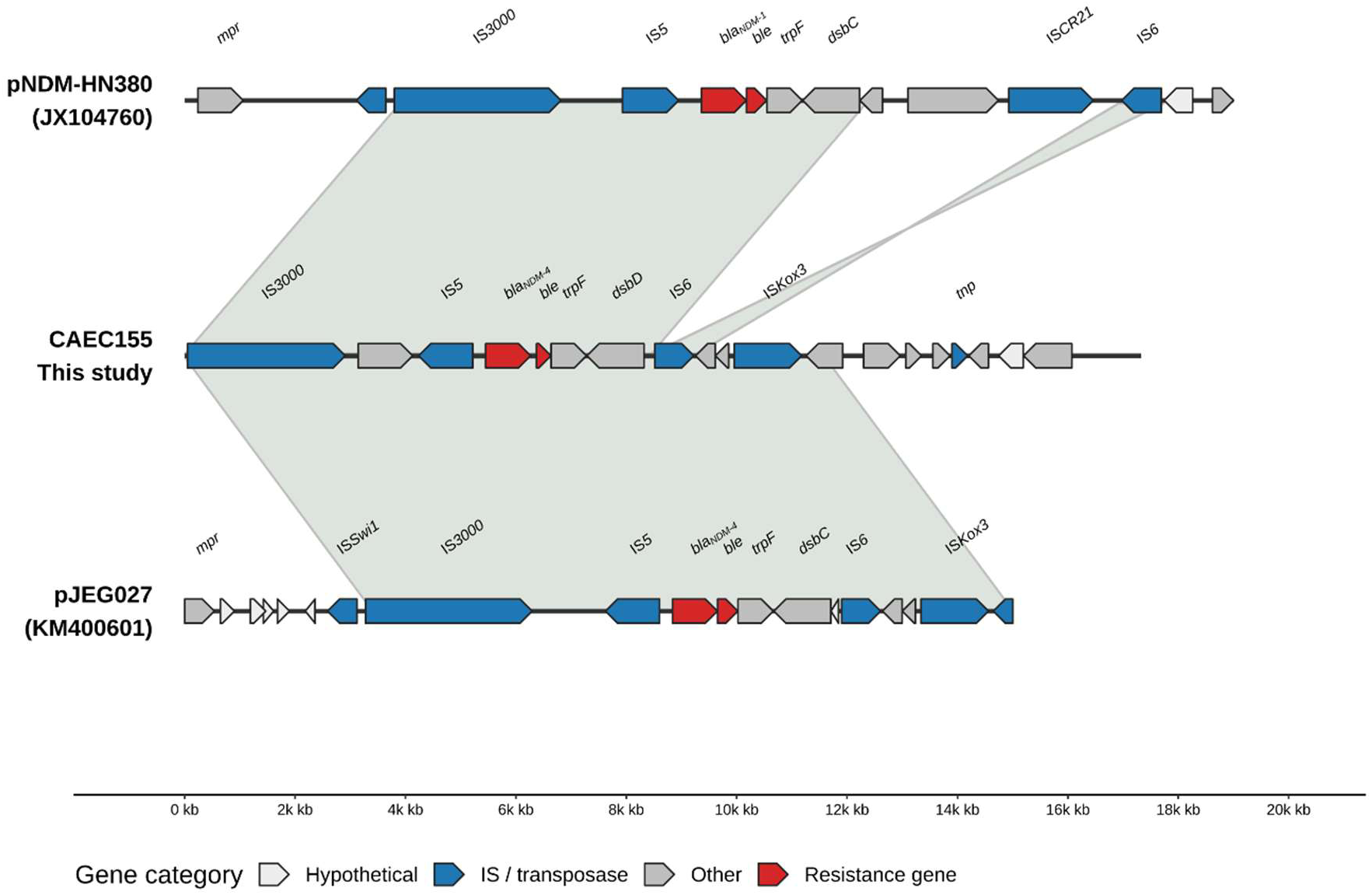
Genetic environment of *bla*_NDM-4_ in CAEC155. Pairwise sequence alignment comparing the NDM-4-bearing IncX3 contig of CAEC155 with reference plasmids pJEG027 (IncX3-*bla*_NDM-4_, accession KM400601) and pNDM-HN380 (IncX3-*bla*_NDM-1_, accession JX104760). Synteny across the IS5-*bla*_NDM_-*ble-trpF-dsbD* cassette are shown. Genes and mobile elements are colored/labeled as per the legend; arrows indicate coding sequence orientation.

### Genetic environment of blaOXA-1207

In CAEC145, *bla*_OXA-1207_ was located on a conjugative IncFII(pCoo)/ColKP3 plasmid. Comparison with reference plasmid pMR3-OXA-181, which carries a complete Tn*6361* transposon bearing *bla*_OXA-181_, showed that CAEC145, EC8729, and EC8533 all carried a ΔTn*6361* element (Figure 2). EC8729 and EC8533 are the two German clinical isolates in which *bla*_OXA-1207_ was originally described. The Tn*6291*-derived block of pMR3-OXA-181 aligned to pIMP-Z1058 at 100% identity over 6,968 bp, and the Tn*2013*-derived block aligned to pKP3-A at 100% identity over 3,022 bp. The three clinical isolates shared a 20,225 bp syntenic block corresponding to part of the Tn*6361* body found in pMR3-OXA-181 at 99.97–99.98% identity but lacked a 1,971 bp internal segment present in the full-length element. Across the full 20,225 bp comparison window, CAEC145 and EC8729 shared 99.98% identity (20,222/20,225 matching bases) and EC8729 and EC8533 shared 99.99% identity (20,224/20,225 matching bases) (Figure 2). The ΔTn*6361* element in CAEC145 was carried on an IncFII(pCoo)/ColKP3 plasmid, whereas EC8729 and EC8533 carried the equivalent element on an IncFIC/ColKP3 plasmid as reported by the original depositors.

**Figure 2.**
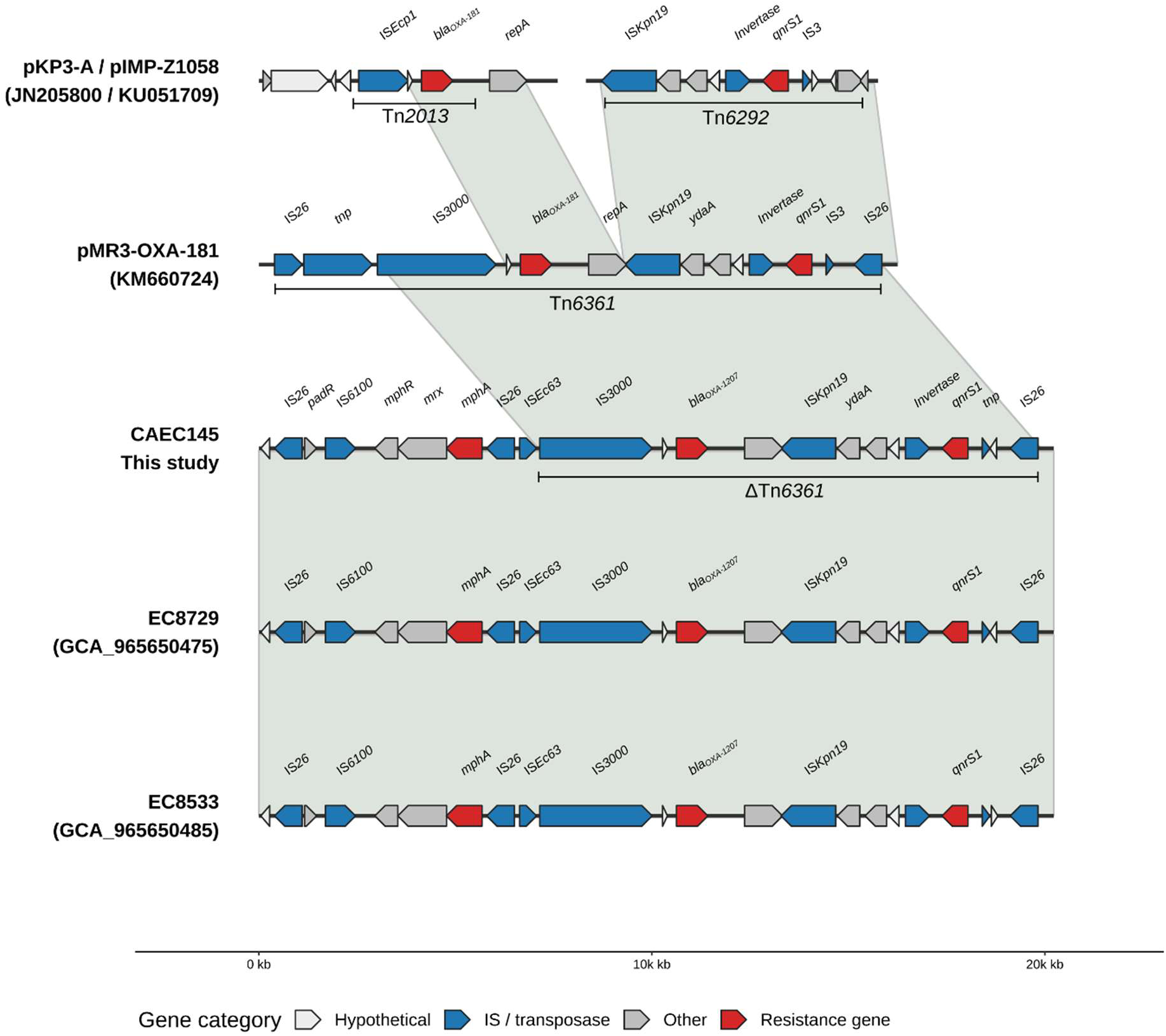
Comparative genetic architecture of the ΔTn*6361* element carrying *bla*_OXA-1207_ in CAEC145. Synteny alignment comparing the OXA-1207-bearing region of CAEC145 with reference plasmid pMR3-OXA-181 (complete Tn*6361* element, accession KM660724) and with the previously reported German isolates EC8729 (GCA_965650475) and EC8533 (GCA_965650485). Additional reference elements pIMP-Z1058 (accession KU051709) and pKP3-A (accession JN205800) are included to delineate the Tn*6291*-and Tn*2013*-derived transposon blocks, respectively. Genes and mobile elements are colored/labeled as per the legend; arrows indicate coding sequence orientation.

### ST361 core-genome phylogeny

Core-genome SNP analysis of 137 ST361 *E. coli* isolates spanning 22 countries (Supplementary Material S1) produced a broadly polyphyletic topology with no single dominant geographic clade. The core alignment covered a mean of 91.6% of the 5.08 Mb reference genome per isolate. The recombination-filtered maximum-likelihood tree was inferred under the TVM+F+ASC+R3 substitution model (log-likelihood −45,056.55; total tree length 0.042 substitutions/site). Lebanese isolates (n=22) resolved across at least three distinct regions of the tree. CAEC145 and CAEC155 were each other’s closest neighbors (28 pairwise SNPs on the recombination-filtered alignment; ML distance 0.0047 substitutions/site), and among all public comparators both were closest to SAMN44683828, a Lebanese clinical isolate collected in 2022 (CAEC155: 63 SNPs, ML distance 0.0109; CAEC145: 76 SNPs, ML distance 0.0132) (Figure 3). Across the 137-isolate set, 82 isolates (59.9%) carried at least one metallo-β-lactamase (78/82 *bla*_NDM-5_), 38 (27.7%) carried an OXA-type carbapenemase, and 4 (2.9%) carried *bla*_KPC-3_. Co-carriage of NDM and OXA determinants was detected in 33 isolates, and one German isolate carried NDM, OXA, and KPC determinants together. *bla*_OXA-1207_ was detected in five isolates: CAEC145 and four isolates from a Singapore hospital surveillance cohort (NOX04, NOX05, NOX08, NOX10; PRJNA1182112; April–May 2024) (31). All four Singapore isolates co-carried *bla*_NDM-5_, whereas CAEC145 carried *bla*_OXA-1207_ as its sole carbapenemase. CAEC145 and the Singapore *bla*_OXA-1207_ carriers did not form an exclusive clade. The *bla*_NDM-4_ allele in CAEC155 was one of only four non-NDM-5 metallo-β-lactamase genes detected in the dataset (the others being *bla*_NDM-1_ in one isolate and unresolved *bla*_NDM_ in two further isolates). Co-carriage of NDM and OXA determinants was not restricted to clinical isolates, having also been detected in environmental wastewater isolates from Kenya (Figure 3).

**Figure 3.**
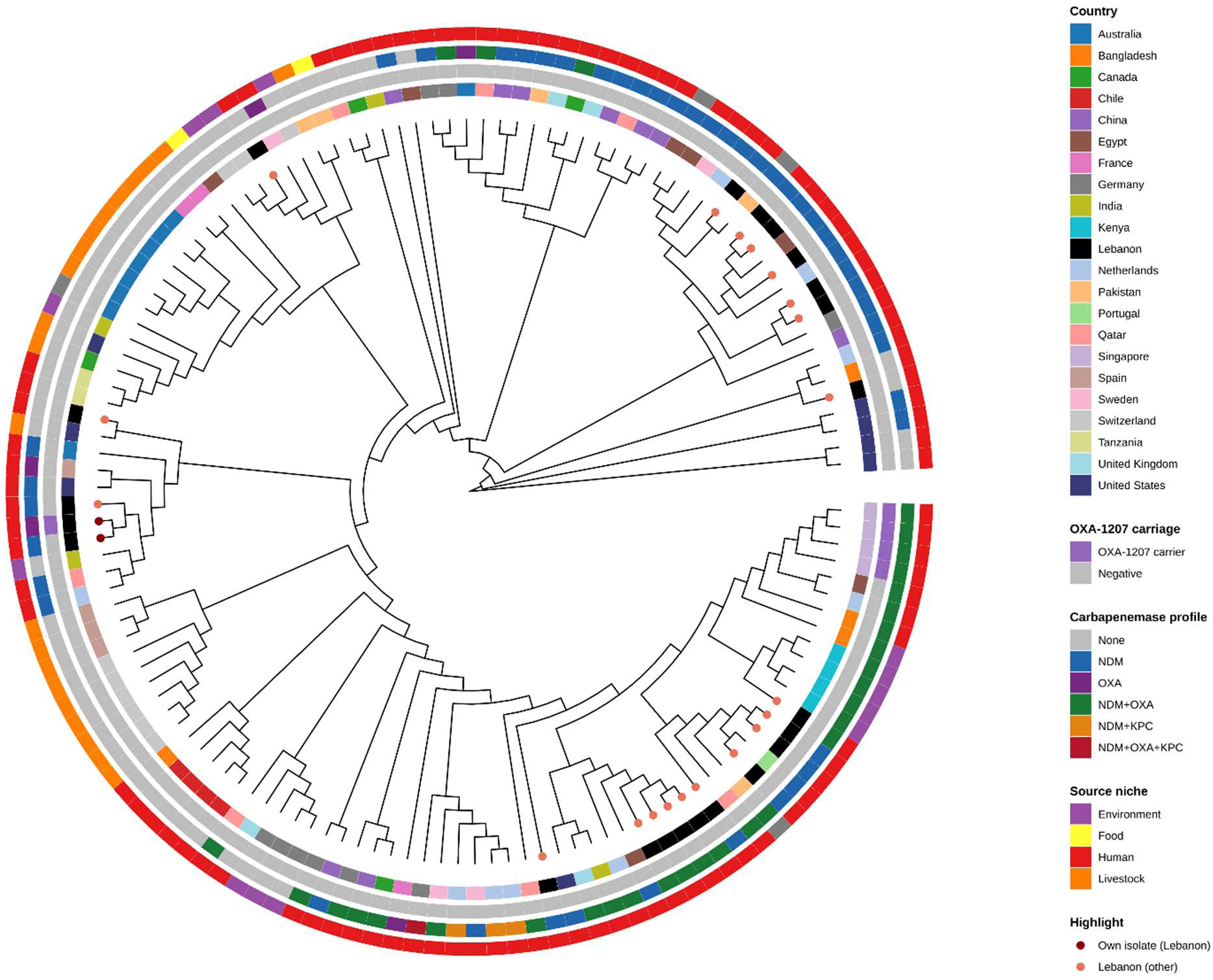
Core-genome phylogeny of 137 global ST361 *Escherichia coli* isolates. Maximum-likelihood phylogeny inferred from a recombination-filtered core-genome SNP alignment against reference genome SAMN53177129 (GCA_057961095.1). The tree is displayed as a circular cladogram annotated by concentric rings indicating, from innermost to outermost: country of origin, OXA-1207 carriage, carbapenemase gene profile (NDM, OXA, KPC, or co-carriage), and source niche (clinical, wastewater, other). CAEC145 and CAEC155 are highlighted; alongside the other Lebanese isolates.

### Antimicrobial susceptibility testing

CAEC145 was susceptible to imipenem (MIC 2 µg/mL) and meropenem (MIC 1 µg/mL) but resistant to ertapenem (Etest MIC >32 µg/mL). CAEC155 was resistant to all three carbapenems (Etest MIC >32 µg/mL each). CAEC145 was susceptible to ceftazidime-avibactam (Etest MIC 2 µg/mL), whereas CAEC155 was resistant (Etest MIC >256 µg/mL). By disk diffusion, both isolates were susceptible to imipenem-relebactam, meropenem-vaborbactam, and cefiderocol, and to amikacin and gentamicin, but resistant to aztreonam, ceftazidime, trimethoprim-sulfamethoxazole, and levofloxacin. Cefepime was reported as resistant for CAEC145 (zone diameter 22 mm, corresponding to susceptible-dose dependent before interpretation) under the CLSI M100 rules applicable to confirmed carbapenemase producers, and CAEC155 showed no zone of inhibition (0 mm, resistant). CAEC145 was susceptible to fosfomycin (27 mm), whereas CAEC155 showed no zone of inhibition (0 mm, resistant). Tigecycline zone diameters were 21 mm and 22 mm for CAEC145 and CAEC155, respectively, reported without categorical interpretation (Table 2).

**Table 2.**
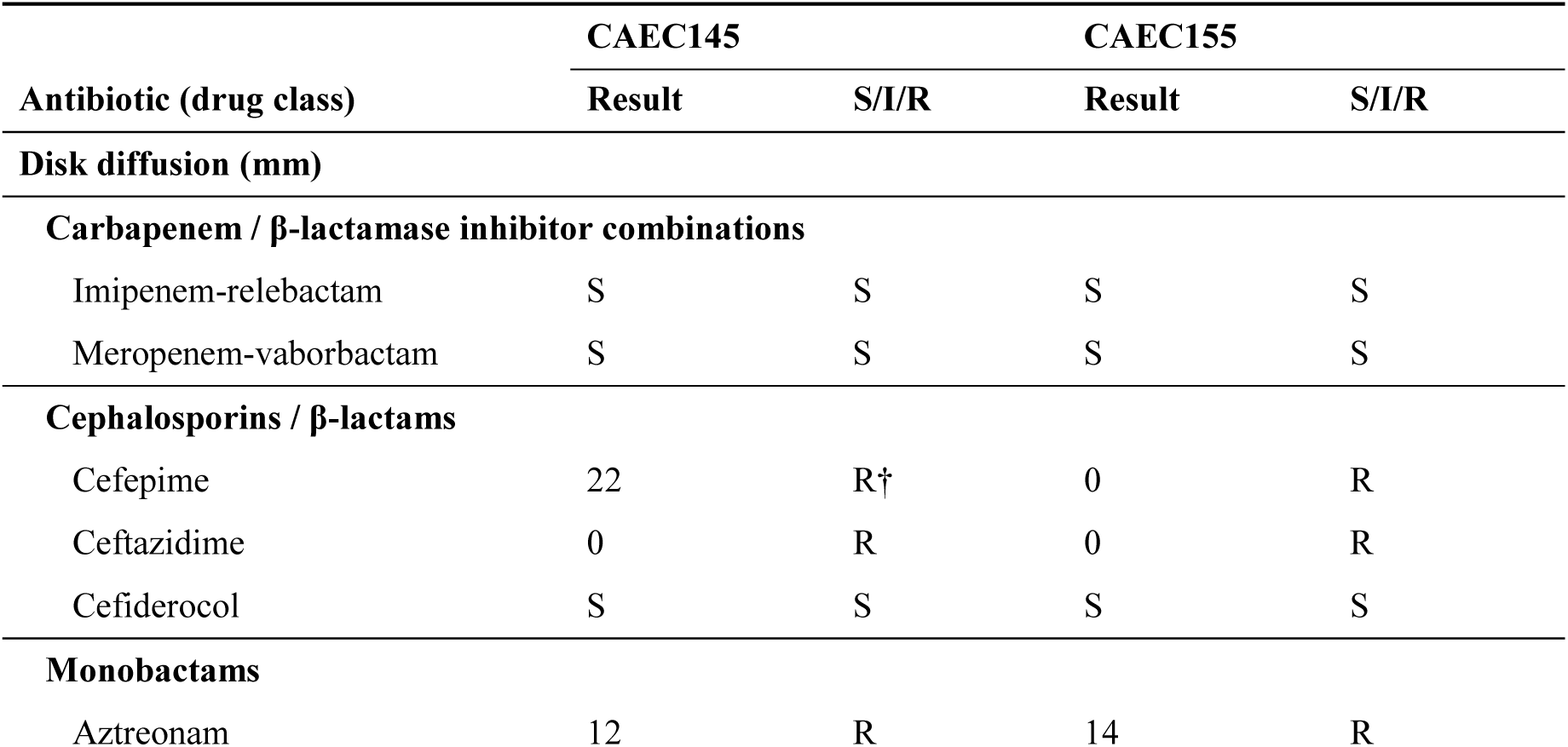
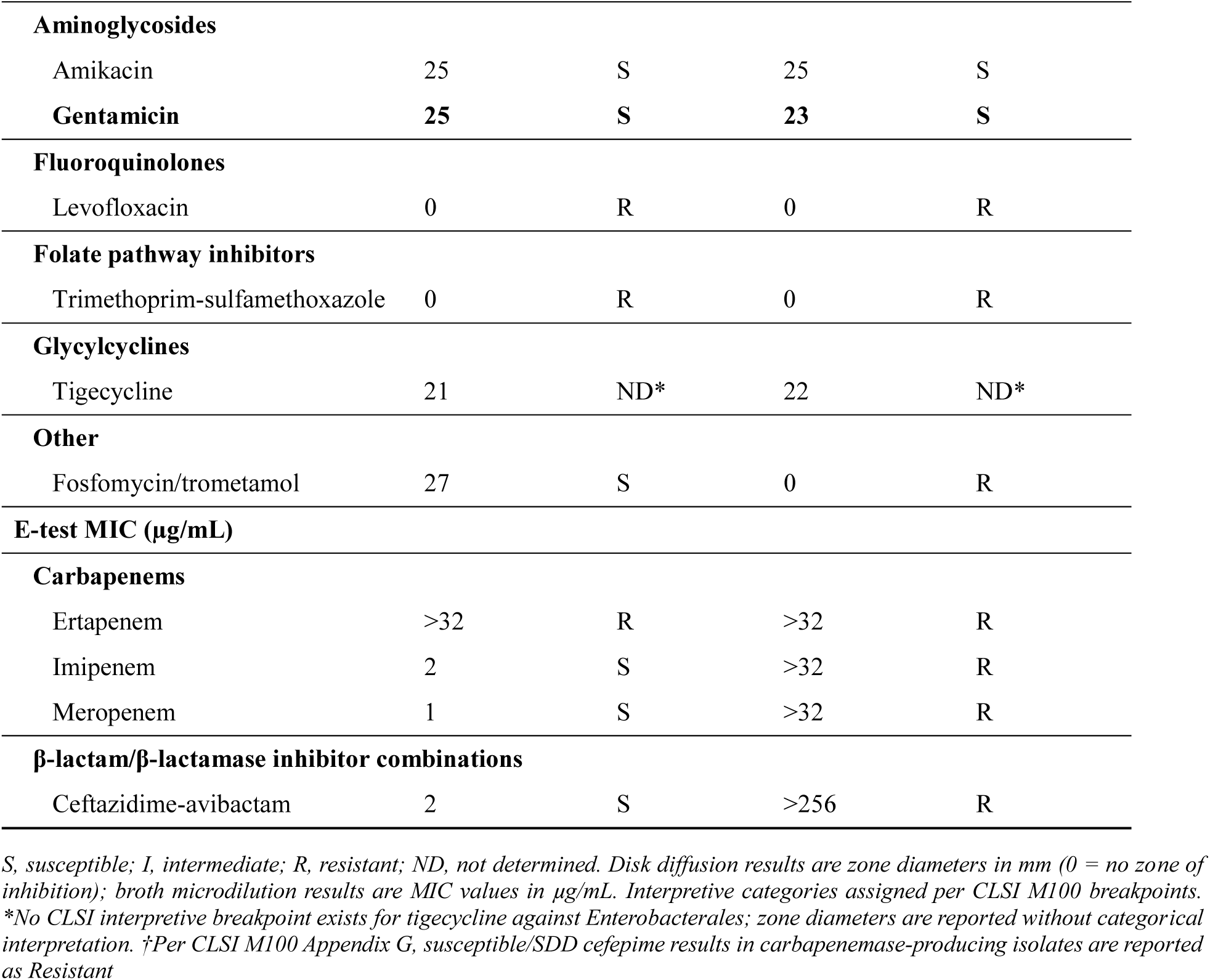
Antimicrobial susceptibility profiles of ST361 *Escherichia coli* isolates CAEC145 and CAEC155 by disk diffusion and E-test.

| Antibiotic (drug class) | CAEC145 |  | CAEC155 |  |
| --- | --- | --- | --- | --- |
|  | Result | S/I/R | Result | S/I/R |
| **Disk diffusion (mm)** |  |  |  |  |
| **Carbapenem / β-lactamase inhibitor combinations** |  |  |  |  |
| Imipenem-relebactam | S | S | S | S |
| Meropenem-vaborbactam | S | S | S | S |
| **Cephalosporins / β-lactams** |  |  |  |  |
| Cefepime | 22 | R† | 0 | R |
| Ceftazidime | 0 | R | 0 | R |
| Cefiderocol | S | S | S | S |
| **Monobactams** |  |  |  |  |
| Aztreonam | 12 | R | 14 | R |
|  | Result | S/I/R | Result | S/I/R |
| **Aminoglycosides** |  |  |  |  |
| Amikacin | 25 | S | 25 | S |
| **Gentamicin** | **25** | **S** | **23** | **S** |
| **Fluoroquinolones** |  |  |  |  |
| Levofloxacin | 0 | R | 0 | R |
| **Folate pathway inhibitors** |  |  |  |  |
| Trimethoprim-sulfamethoxazole | 0 | R | 0 | R |
| **Glycylcyclines** |  |  |  |  |
| Tigecycline | 21 | ND* | 22 | ND* |
| **Other** |  |  |  |  |
| Fosfomycin/trometamol | 27 | S | 0 | R |
| **E-test MIC (µg/mL)** |  |  |  |  |
| **Carbapenems** |  |  |  |  |
| Ertapenem | >32 | R | >32 | R |
| Imipenem | 2 | S | >32 | R |
| Meropenem | 1 | S | >32 | R |
| **β-lactam/β-lactamase inhibitor combinations** |  |  |  |  |
| Ceftazidime-avibactam | 2 | S | >256 | R |

S, susceptible; I, intermediate; R, resistant; ND, not determined. Disk diffusion results are zone diameters in mm (0 = no zone of inhibition); broth microdilution results are MIC values in µg/mL. Interpretive categories assigned per CLSI M100 breakpoints.
\*No CLSI interpretive breakpoint exists for tigecycline against Enterobacterales; zone diameters are reported without categorical interpretation. †Per CLSI M100 Appendix G, susceptible/SDD cefepime results in carbapenemase-producing isolates are reported as Resistant

## Discussion

The central finding of this study is that CAEC145 and CAEC155, recovered from the same patient during a single sequential infection episode and separated by only 28 core-genome SNPs, differ in the class of carbapenemase they carry rather than in its allelic variant. The OXA-48-family *bla*_OXA-1207_ carried by CAEC145 was entirely absent from CAEC155, replaced by *bla*_NDM-4_, a class B1 metallo-β-lactamase. Within-host evolution of carbapenem-resistant *E. coli* has previously been documented mainly as porin loss combined with pre-existing β-lactamase expression (32), or as retention of a single resistant clone across infection episodes (5). In a recent longitudinal study of within-host CREc populations comprising 20 clonal isolate groups from patients with recurrent or multi-site infection, no acquisition or loss of carbapenemase genes was observed in any patient (5). By contrast, the complete within-host succession documented here, from one carbapenemase class to an enzymatically unrelated one and confirmed by hybrid whole-genome sequencing, represents a clear deviation from the pattern of stable within-host carbapenemase content. To our knowledge, it is also the first such event reported for *bla*_OXA-1207_.

The phenotypic data illustrate the direct clinical consequences of the carbapenemase class switch. The OXA-48-family background of CAEC145 preserved susceptibility to ceftazidime-avibactam, whose avibactam component effectively inhibits these serine β-lactamases (6), alongside imipenem-relebactam, meropenem-vaborbactam, and cefiderocol, leaving multiple treatment options available. The acquisition of *bla*_NDM-4_ in CAEC155 abolished ceftazidime-avibactam susceptibility and conferred high-level resistance to imipenem and meropenem, the two carbapenems that had remained active against CAEC145. The concurrent loss of fosfomycin susceptibility in CAEC155, attributable to acquisition of *fosA4*, narrowed the remaining options further. The agents that remained active against both isolates (imipenem-relebactam, meropenem-vaborbactam, cefiderocol, amikacin, and gentamicin) define the shared therapeutic options. However, the transition from CAEC145 to CAEC155 effectively eliminated the only carbapenemase-targeting β-lactam and the two carbapenems clinicians rely on most, a consequence that a resistance profile from either isolate alone would not have revealed.

The plasmid configuration is consistent with sequential acquisition and displacement under sustained carbapenem selection, rather than in-place evolution of a single gene. The *bla*_OXA-1207_ plasmid of CAEC145 and the *bla*_NDM-4_ replicon of CAEC155 belong to different Inc groups, so neither replication nor partitioning incompatibility would have prevented their coexistence or the replacement of one by the other (15). The IncX3-type *bla*_NDM-4_ cassette in CAEC155 was nearly identical (99.78% identity across the IS*5*-*bla*_NDM_-*ble*-*trpF*-*dsbD* unit) to pJEG027, an IncX3-*bla*_NDM-4_ plasmid originally recovered from a *K. pneumoniae* isolate from an Australian patient (33). It differed from the prototype IncX3-*bla*_NDM-1_ plasmid pNDM-HN380 only at the expected NDM substitution, indicating that CAEC155 acquired a typical, largely unrearranged member of this globally disseminated lineage rather than generating a locally novel construct. IncX3 plasmids are a well-established vehicle for global *bla*_NDM_ dissemination, characterized by high conjugative transfer efficiency, minimal fitness burden on their Enterobacterales host, and a narrow but clinically dominant host range (18, 34). Under the selective pressure of carbapenem therapy during a relapsing infection, acquiring this stable, highly transmissible plasmid likely allowed the strain to outcompete or displace the costlier *bla*_OXA-1207_ plasmid within the host. *bla*_OXA-1207_ is itself a recently emerged variant of the *bla*_OXA-181_ family with a widening geographic distribution. First described in mid-2025 in two unrelated German ST405 and ST4405 isolates on a plasmid whose conjugation frequency (mean 1.2 × 10⁻¹) substantially exceeded rates typical of established OXA-48-family plasmids (12), it was subsequently reported in an ST617 isolate from a pediatric patient in Qatar (13). Within the 137-genome ST361 phylogeny, *bla*_OXA-1207_ appeared independently in Lebanon (CAEC145) and Singapore on non-monophyletic backgrounds. In four concurrent Singapore ST361 isolates it was co-carried with *bla*_NDM-5_, whereas in CAEC145 it was the sole carbapenemase. The ΔTn*6361* structure linked CAEC145 to the original German isolates at up to 99.99% identity across a 20 kb window, evidence of a shared mosaic origin. The flanking backbone, however, diverged: *bla*_OXA-1207_ was carried on an IncFII(pCoo)/ColKP3 plasmid in CAEC145 and on an IncFIC/ColKP3 plasmid in the German isolates. This pattern is consistent with a conserved transposon core recombining onto whichever compatible replicon is locally available.

The phylogenomic context reinforces that this isolate pair is not an epidemiologically isolated event. Both CAEC145 and CAEC155 were more closely related to a Lebanese clinical isolate from 2022 than to any other public ST361 genome. This implies local circulation for at least three years before sampling, though a single comparator genome cannot rule out recent, unsampled importation. More broadly, carbapenemase carriage across the 137-genome ST361 dataset was widespread and scattered across unrelated clades, spanning clinical and wastewater sources and including isolates from Kenya. This distribution is more consistent with repeated, independent horizontal acquisition within an already globally disseminated lineage, rather than clonal expansion of a single resistant strain (35).

These findings document, for the first time to our knowledge, a within-host succession from an OXA-48-like to an NDM-type carbapenemase over the course of a single sequential *E. coli* infection. The switch appears driven by plasmid-level displacement rather than in-place gene evolution, set within an endemic, phylogenetically diverse, and still-expanding ST361 lineage. This argues for genomic, rather than purely phenotypic, surveillance of nosocomial CRE outbreaks (36). A carbapenemase class switch of this kind carries direct implications for definitive antibiotic selection that a static resistance profile from a single isolate would not reveal.

### Limitations

As a single-patient, two-isolate case report, this study establishes that a within-host carbapenemase class switch can occur but cannot quantify how often it does; the true frequency requires systematic paired-isolate sequencing across larger sequential-infection cohorts. Several further constraints apply. Conjugative transfer of the *bla*_NDM-4_ and *bla*_OXA-1207_ plasmids was predicted from replicon content rather than demonstrated by mating assay, and the fitness cost invoked to explain displacement was not measured. With only two isolates available, we could not establish whether *bla*_OXA-1207_ was lost before, during, or after *bla*_NDM-4_ acquisition, nor exclude an unsampled intermediate population carrying both determinants.

## Data Availability

The draft genome sequences of strains CAEC145 and CAEC155 have been deposited at the National Center for Biotechnology Information (NCBI) under BioProject PRJNA1413914. The GenBank accession numbers are JBTYJV000000000 for CAEC145 and JBTYJW000000000 for CAEC155.

## Declaration of generative AI and AI-assisted technologies in the manuscript preparation process

During the preparation of this work, the authors used ChatGPT and Claude to improve language and readability. After using these tools, the authors reviewed and edited the content as needed and take full responsibility for the content of the published article.

## Supplemental Material

Supplemental Material 1. Metadata and carbapenemase gene profiles of the 137 ST361 *Escherichia coli* isolates used in core-genome phylogenetic analysis

Supplemental Material 2. Complete AMR genes list of CAEC145 and CAEC155

Supplemental material 3. Complete virulence genes list of CAEC145 and CAEC155

